# Computationally guided design of a metastasis-on-a-chip platform for quantitative evaluation of chemotactic cues in developmental cancers

**DOI:** 10.64898/2026.07.25.740695

**Authors:** Catherine Murphy, Luka Jarc, Alessia Cadavere, Emilia Cioffi, Maider Badiola-Mateos, David Fernandez, Nathalia Gomez-Jimenez, Jaume Mora, Josep Samitier, Aranzazu Villasante

## Abstract

Metastatic dissemination is initiated by tumor cells interpreting spatially organized biochemical and biophysical cues that remain difficult to reproduce using conventional migration assays. Here, we developed a computationally guided metastasis-on-a-chip (MET-on-a-chip) platform based on the concept of the Minimally Functional Unit (MFU), in which only the biological components required to answer a defined experimental question are incorporated. The platform consists of two independent culture chambers connected through an array of confined microchannels that permits diffusion of soluble factors while constraining tumor cell migration. Rather than relying on empirical optimization, finite-element COMSOL simulations were first used to predict molecular transport, define growth factor loading conditions, identify biologically relevant exposure regions, and guide the rational design of the microfluidic assay.

Computational predictions were experimentally validated using 70-kDa FITC-dextran diffusion and VEGF release studies, confirming the formation of stable spatial concentration gradients across the microfluidic platform. The simulations further demonstrated that both growth factor loading and cell positioning relative to the predicted gradients critically influenced assay performance, leading to the optimization of the platform through spatial reconfiguration of the tumor compartment.

Using the optimized configuration, we compared the migratory responses of neuroblastoma, Ewing sarcoma, and osteosarcoma cells to vascular (VEGF-A165) and lymphatic (VEGF-C) chemotactic cues. VEGF-C significantly increased migration through the microchannel array in Ewing sarcoma and osteosarcoma cells, whereas VEGF-A165 produced no significant effect. In contrast, neuroblastoma cells exhibited minimal migration under either condition, revealing tumor-specific differences in responsiveness to VEGF signaling.

Together, these findings establish a computationally guided workflow for the rational design of metastasis-on-a-chip assays, in which predictive modeling informs experimental design before biological validation. By substantially reducing empirical trial-and-error while enabling quantitative control over growth factor exposure, this strategy provides a robust framework for developing minimally functional microphysiological systems capable of dissecting individual steps of the metastatic cascade under experimentally defined conditions.

**Translational Impact Statement:** Metastatic dissemination remains one of the greatest clinical challenges in pediatric oncology, yet experimental models capable of quantitatively evaluating early migratory events remain limited. The computationally guided MET-on-a-Chip workflow presented here provides a human-relevant platform in which soluble microenvironmental cues can be systematically investigated under controlled and predictive conditions. Although demonstrated here using VEGF-A165 and VEGF-C, the platform can be readily adapted to study virtually any chemotactic factor, cytokine, extracellular vesicle population, or therapeutic candidate involved in metastatic dissemination.

The modular MFU design allows biological complexity to be incorporated progressively as dictated by the scientific question, providing a flexible framework for future applications.

In the longer term, this workflow could be combined with patient-derived tumor cells, organoids, or biopsy material to investigate patient-specific metastatic behavior and evaluate anti-metastatic therapeutic strategies in a personalized setting. Beyond identifying pro-migratory signaling pathways, the platform may serve as a preclinical tool to prioritize compounds capable of preventing tumor cell dissemination before evaluation in more complex animal models or clinical studies.

## Introduction

Metastasis remains the leading cause of death in children with solid tumors, including neuroblastoma (NB), Ewing sarcoma (ES), and osteosarcoma (OS). Although considerable advances have improved the treatment of localized disease, the prognosis of patients with metastatic tumors has changed only modestly over the past decades^1–4^. One of the major reasons for this limited progress is that the earliest stages of metastatic dissemination remain poorly understood, particularly the mechanisms by which tumor cells interpret and respond to spatially organized signals within their microenvironment^5^.

Tumor dissemination is not a random process. Before entering the circulation and colonizing distant organs, cancer cells continuously integrate biochemical and biophysical information provided by the surrounding tissue. Soluble growth factors, extracellular matrix composition, tissue architecture, mechanical confinement, vascular organization, and stromal cells collectively shape migratory behavior^6^. Importantly, these signals are neither homogeneous nor static. Instead, they exist as dynamic spatial gradients that evolve over time and differ substantially across tissues. Reproducing these gradients experimentally remains one of the major challenges in metastasis research^7–9^.

Traditional migration assays, including Boyden chambers, Transwell systems, matrigel or collagen invasion assays, and other endpoint-based approaches, have been instrumental in identifying molecules involved in tumor cell motility^10–12^. However, these methods provide limited control over the spatial and temporal organization of chemotactic cues. Concentration gradients are poorly defined, progressively dissipate during the experiment, and cannot be directly related to the concentration actually experienced by individual cells. Furthermore, endpoint measurements make it difficult to distinguish directed chemotaxis from increased random motility (chemokinesis), limiting mechanistic interpretation of migratory responses^10–12^.

Microfluidic organ-on-a-chip technologies have emerged as a powerful alternative because they enable the generation of defined tissue architectures together with controlled molecular gradients, mechanical constraints, and multicellular interactions^13,14^. Rather than attempting to reproduce the entire metastatic cascade, these systems make it possible to isolate individual biological processes under experimentally controlled conditions. This reductionist engineering approach represents one strategy for developing human microphysiological systems in which biological complexity is introduced only when required to reproduce the function under investigation^13,14^.

In our view, this concept can be formalized as the Minimally Functional Unit (MFU): the simplest experimental system capable of reproducing the biological function required to answer a specific scientific question. The objective is therefore not to maximize biological complexity, but to include only those components that are functionally necessary for the phenomenon being investigated. Additional biological elements should be incorporated only when they provide mechanistic information that cannot be obtained using a simpler system^15^. This engineering philosophy underpins the present work. Besides facilitating experimental interpretation, it improves reproducibility and enables systematic interrogation of individual microenvironmental cues while providing a modular framework that can progressively evolve toward more complex human-relevant models.

Despite the rapid development of organ-on-a-chip technologies, the design of new microphysiological systems still relies largely on empirical optimization. Device geometry, hydrogel composition, loading concentrations, cell positioning, and experimental timing are commonly established through repeated trial-and-error iterations, requiring multiple fabrication cycles and consuming considerable amounts of time, reagents, and valuable biological material before biologically meaningful conditions are achieved. Computational modeling offers an attractive alternative by allowing molecular transport, gradient formation, and spatial exposure profiles to be predicted before experiments are performed. Rather than serving only to explain experimental observations retrospectively, finite-element modeling can become an integral component of assay design, enabling rational optimization of experimental conditions while substantially reducing empirical testing^16,17^.

This concept is particularly relevant for members of the vascular endothelial growth factor (VEGF) family, which regulate angiogenesis, lymphangiogenesis, vascular permeability, and tissue remodeling^18,19^. VEGF-A165 primarily signals through VEGFR2 (KDR) and is the principal driver of angiogenic sprouting, whereas VEGF-C preferentially activates VEGFR3 (FLT4) and plays a central role in lymphatic vessel development and metastatic dissemination. Beyond their effects on endothelial cells, accumulating evidence indicates that both ligands can directly influence tumor cell migration through autocrine and paracrine mechanisms, although these responses appear to depend strongly on tumor type and biological context^18,19^. In pediatric cancers, VEGF signaling has been associated with tumor progression, angiogenesis, metastatic dissemination, and adverse clinical outcome in neuroblastoma, Ewing sarcoma, and osteosarcoma, supporting the biological relevance of this signaling axis in developmental tumors^20^.

Importantly, the concentrations of VEGF experienced by cells in vivo are considerably lower than those commonly loaded into experimental migration assays. A comprehensive meta-analysis estimated VEGF-A concentrations between approximately 13.3 and 39.4 pg/mL within the tumor interstitium, compared with approximately 74 pg/mL in serum and 328 pg/mL in plasma from cancer patients, illustrating the steep concentration gradients that naturally exist between the tumor microenvironment and the circulation^21^. Consequently, the concentration initially loaded into a hydrogel or microfluidic device should not be assumed to represent the concentration ultimately experienced by responding cells after diffusion through extracellular matrices, reservoirs, and connecting microchannels. Instead, local exposure depends on molecular transport, device geometry, and cell positioning, emphasizing the need for predictive computational approaches capable of quantitatively defining these gradients before biological experimentation^16,17^.

Here, we present a computationally guided MET-on-a-Chip platform developed according to the MFU concept to investigate VEGF-dependent migration in pediatric solid tumors. Rather than recreating the entire metastatic niche, we designed the minimal microphysiological system required to address a single biological question: whether vascular (VEGF-A165) and lymphatic (VEGF-C) cues are sufficient to induce directional migration in neuroblastoma, Ewing sarcoma, and osteosarcoma cells. Finite-element simulations were first used to predict molecular transport, define growth factor loading conditions, identify biologically relevant exposure regions, and optimize the experimental configuration before fabrication and biological validation. This predictive workflow establishes a rational strategy for designing metastasis-on-a-chip assays while revealing tumor-specific responses to vascular and lymphatic chemotactic cues.

## Materials and Methods

### Analysis of VEGF receptor expression in developmental solid tumors

To identify clinically relevant chemotactic factors for evaluation in the MET-on-a-Chip platform, the expression of the VEGF receptors KDR (VEGFR2) and FLT4 (VEGFR3) was analyzed using publicly available pediatric cancer gene expression datasets available through the R2 Genomics Analysis and Visualization Platform (https://r2.amc.nl).

Datasets corresponding to neuroblastoma, Ewing sarcoma and osteosarcoma were selected from the R2 MegaSampler collection. Gene expression values were obtained using the corresponding Affymetrix probe sets (KDR: 203934_at; FLT4: 229902_at) and are displayed as log2-transformed normalized expression values. Box-and-whisker plots were generated directly using the R2 platform, allowing comparison of receptor expression across tumor entities and independent patient cohorts.

### MET-on-a-Chip design and fabrication

The MET-on-a-Chip platform was designed using AutoCAD (Autodesk, USA) and fabricated by combining standard photolithography and PDMS soft lithography techniques. The device comprises two independent culture chambers connected to two intermediate reservoirs through access microchannels. The reservoirs are interconnected by an array of parallel microchannels that establishes the diffusion interface between both sides of the device. The complete design was transferred to two independent acetate photomasks corresponding to the first (microchannels) and second (chambers and reservoirs) lithographic layers (CAD/Art Services Inc., OR, USA).

Master molds were fabricated on 4-inch silicon wafers under clean-room conditions using a two-step photolithographic process. Before photoresist deposition, wafers were cleaned by oxygen plasma treatment (20 min, 7.2 W, 0.2 Torr; Harrick Plasma PDC-002-CE). The first lithographic layer, corresponding to the migration microchannels, was fabricated using SU-8 2010 photoresist spin-coated to obtain a nominal thickness of 15 μm. After soft baking, the wafer was exposed through the corresponding acetate mask using a manual mask aligner (SÜSS MicroTec MJB4), followed by post-exposure baking.

A second lithographic step was subsequently performed using SU-8 2100 photoresist to generate the chambers, reservoirs and access microchannels. Alignment between both layers was achieved using dedicated alignment marks incorporated into the photomasks. Following exposure, the masters were developed in SU-8 developer, rinsed with isopropanol, hard baked and silanized under vacuum using trichloro(1H,1H,2H,2H-perfluorooctyl)silane vapor before replica molding.

PDMS replicas were obtained by pouring a degassed 10:1 mixture of Sylgard 184 silicone elastomer base and curing agent (Dow Corning) onto the silanized masters and curing overnight at 65°C. After curing, PDMS replicas were removed from the master, reservoirs were opened using a 10-mm biopsy punch and culture chambers were generated using a custom-made aluminum punch matching the chamber geometry. Devices were then permanently bonded to 24 × 60 mm glass coverslips by oxygen plasma activation.

Immediately before use, bonded devices were washed with ethanol, dried, treated with UV-ozone to restore surface hydrophilicity, filled with sterile Milli-Q water and sterilized under UV light until cell seeding.

All experiments were performed using this MET-on-a-Chip device, which incorporates a 1-mm separation between culture chambers to improve fabrication reproducibility and facilitate long-range migration experiments.

### MET-on-a-Chip characterization

The geometry of the fabricated PDMS devices was experimentally characterized prior to biological experiments and computational modeling. Lateral dimensions, including chamber diameters, reservoir dimensions, access microchannels, and the migration microchannel array, were measured from optical micrographs acquired using an inverted microscope (Olympus IX71, Olympus, Japan) and quantified using Fiji/ImageJ (n = 5 devices).

The height of the different SU-8 structures and PDMS replicas was measured using a DektakXT stylus profilometer (Bruker, USA) equipped with a 12.5-µm radius stylus operated with an applied force of 3 mg. Step-height measurements were performed according to the manufacturer’s operating recommendations.

Because the migration microchannels (15 µm wide) approach the lateral dimensions of the profilometer stylus, their depth was additionally verified using a Wyko NT1100 optical interferometer (Veeco Instruments, USA) operated in Vertical Scanning Interferometry (VSI) mode. Samples were levelled before acquisition, and channel depths were determined from two-dimensional optical profiles generated using Vision32 software.

To evaluate the robustness of the fabrication process, microchannel depth was additionally quantified in PDMS devices polymerized under different curing conditions.

### Cell culture

Human neuroblastoma SK-N-BE(2), SK-N-MC (Ewing sarcoma) and U2OS (osteosarcoma) cell lines were obtained from the Pediatric Cancer Center Barcelona, Hospital Sant Joan de Déu (Barcelona, Spain). The SK-N-BE(2)-Tomato cell line was generated by stable tdTomato transduction by Dr. Raquel Bernard in the laboratory of Dr. Manuel Serrano (IRB Barcelona).

Cells were cultured in RPMI-1640 medium supplemented with 10% fetal bovine serum (FBS), 1% penicillin–streptomycin and maintained at 37°C in a humidified atmosphere containing 5% CO□. Culture medium was replaced every 2–3 days and cells were routinely passaged upon reaching approximately 80% confluence.

### MET-on-a-Chip preparation and hydrogel loading

Microfluidic connectivity and microchannel integrity were verified by monitoring the diffusion of colored Medium 199 through the microchannel array prior to each experiment. Devices showing incomplete microfluidic connectivity or damaged microchannels were discarded. Successfully validated devices were subsequently activated by UV-ozone treatment for 15 min, sterilized under UV light for an additional 15 min and coated with gelatin for 1 h at room temperature to promote hydrogel adhesion. Following removal of the gelatin solution, devices were washed with PBS and hydrated before hydrogel loading.

Chamber 2 was prepared one day before cell seeding by loading 200 µL of a neutralized collagen type I hydrogel (Corning #354249) at a final collagen concentration of 5 mg mL□¹, supplemented with either recombinant human VEGF-A165 (3.5 µg mL□¹; Gibco, Cat. No. 100-20-10UG) or recombinant human VEGF-C (5 µg mL□¹; Gibco, Cat. No. 100-20CD-20UG). Control devices received an identical collagen hydrogel without growth factor supplementation. Following loading, devices were maintained overnight at 4 °C to allow complete hydrogel polymerization.

On the following day, cancer cells embedded in a collagen I–fibrin hydrogel were loaded either into Chamber 1 (configuration 1) or Reservoir 1 (configuration 2). Unless otherwise indicated, cancer cells were labeled for 1 h at 37 °C with CellTracker fluorescent probes (Thermo Fisher Scientific, Waltham, MA, USA) according to the manufacturer’s instructions before encapsulation in the hydrogel. SK-N-BE(2)-Tomato cells were used without additional staining owing to their stable tdTomato expression.

For configuration 1, the hydrogel consisted of collagen type I (Corning #354249; final concentration 4 mg mL□¹), fibrinogen (Sigma-Aldrich #F8630; final concentration 10 mg mL□¹), thrombin (Sigma-Aldrich #T4648; final concentration 2.5 U mL□¹), aprotinin (Sigma-Aldrich #A6106; final concentration 80 µg mL□¹), Medium 199 (Sigma-Aldrich #M9163-), HEPES (10 mM) and NaHCO□ (15 mM). The pH was adjusted to approximately 7.5 before cell incorporation. Cancer cells were resuspended in the thrombin solution and mixed with the pre-hydrogel immediately before loading to obtain a final density of 1.5 × 10□ cells mL□¹. Two hundred microlitres of the cell-laden hydrogel were introduced into Chamber 1 and allowed to polymerize on a 37 °C hotplate for approximately 90 s. Finally, complete RPMI-1640 medium supplemented with 10% fetal bovine serum and 1% penicillin–streptomycin was added to both chambers (200 µL) and reservoirs (100 µL), and the devices were incubated until imaging.

In configuration 2, a 50-µL collagen I–fibrin hydrogel containing 112,500 cancer cells (2.25 × 10□ cells mL□¹) was polymerized within Reservoir 1, leaving sufficient space for 50 µL of culture medium above the hydrogel. The hydrogel composition and polymerization conditions were otherwise identical to those used for Chamber 1 seeding. This configuration established a unidirectional VEGF gradient from Chamber 2 toward Chamber 1 through the microchannel array.

### Computational modeling

#### Computational modeling of VEGF-A165 diffusion

Three-dimensional diffusion simulations were performed using COMSOL Multiphysics® version 6.2 (COMSOL Inc., Burlington, MA, USA) to predict the establishment of VEGF-A165 and VEGF-C concentration gradients within the MET-on-a-Chip platform before experimental validation. The computational model was developed using the experimentally measured dimensions of the fabricated MET-on-a-Chip device, including culture chambers, intermediate reservoirs, connecting channels and the array of 125 parallel microchannels described above.

The computational geometry reproduced the complete architecture of the microfluidic platform. Chamber 2 contained the VEGF-loaded collagen hydrogel, whereas all remaining domains were modeled as culture medium. The collagen hydrogel was represented as a porous material, while the surrounding domains were assigned the physical properties of water-based culture medium.

VEGF transport was simulated using the Transport of Diluted Species module in COMSOL Multiphysics®. Transport of diluted species was described by the mass conservation equation:

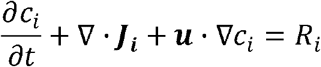

with

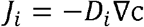

where

- *c_i_*: concentration of species *i* (mol m□³)
- *n_i_*: diffusion coefficient (m² s□¹)
- *R_i_*: reaction term (mol m□³ s□¹)
- ***u***: mass-averaged velocity vector (m s□¹)
- ***J_i_***: diffusive flux (mol m□² s□¹)

Because all simulations reproduced static culture conditions, convective transport was neglected. Accordingly, VEGF transport within the device was modeled as diffusion coupled to first-order degradation according to

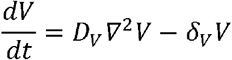

where

- *D_V_*: diffusion coefficient
- *o_V_*: degradation rate

The same computational framework, governing equations and device geometry were used for both VEGF-A165 and VEGF-C simulations. Independent simulations were nevertheless performed for each growth factor to determine the minimum loading concentration capable of generating a measurable and sustained concentration gradient throughout the device under the corresponding experimental conditions. Thus, the simulations were used to calibrate the source concentration incorporated into the collagen hydrogel rather than to compare the intrinsic diffusion properties of the two molecules.

Time-dependent simulations were performed between 0 and 24 h for VEGF-A165 and between 0 and 48 h for VEGF-C, matching the duration of the corresponding migration assays. Several initial loading concentrations were evaluated for each growth factor through parametric analyses before selecting the experimental conditions used for biological validation.

To quantify VEGF transport throughout the device, concentration profiles were extracted along predefined three-dimensional Cut Lines positioned at successive regions of the microfluidic platform. For VEGF-A165, concentration profiles were analyzed in the channel connecting Chamber 2 to Reservoir 2, Reservoir 2, Reservoir 1 and Chamber 1. For VEGF-C, the analysis focused on three regions of interest corresponding to Channel 2, the microchannel array connecting both reservoirs and Channel 1 adjacent to Chamber 1, allowing quantitative assessment of growth factor attenuation along the complete diffusion path.

#### Quantification of VEGF release from collagen hydrogels

To experimentally validate the growth factor loading conditions predicted by the computational models, the release of VEGF-A165 and VEGF-C from collagen hydrogels was quantified by enzyme-linked immunosorbent assay (ELISA). Acellular collagen hydrogels containing the corresponding growth factor at the loading concentrations selected from the COMSOL simulations (3.5 μg/mL VEGF-A165 or 5 μg/mL VEGF-C) were prepared under the same conditions used for the migration experiments.

Hydrogels were incubated for 24 h either under gel polymerization conditions (PBS, 4°C) or under standard cell culture conditions (RPMI, 37°C). After incubation, the complete supernatant (600 μL) was collected from each hydrogel for analysis.

VEGF-A165 and VEGF-C concentrations were quantified using commercially available human ELISA kits (RayBiotech, Peachtree Corners, GA, USA; Catalog Nos. ELH-VEGF and ELH-VEGFC, respectively), following the manufacturer’s instructions. Briefly, standards and samples were added to antibody-coated 96-well plates and incubated for 2.5 h at room temperature, followed by incubation with biotinylated detection antibody (1 h), HRP-conjugated streptavidin (45 min), and TMB substrate (30 min). The reaction was stopped with sulfuric acid and absorbance was measured at 450 nm using a microplate reader. Concentrations were calculated from standard curves generated for each assay according to the manufacturer’s protocol.

Four independent collagen hydrogels were analyzed for each condition (n = 4), and results are presented as mean ± standard deviation.

#### Experimental validation of COMSOL-predicted diffusion gradients

To experimentally validate the diffusion gradients predicted by the COMSOL simulations, the MET-on-a-Chip devices described above were prepared following the same fabrication, coating and hydrogel-loading procedure. The only modification consisted of replacing recombinant VEGF with 70-kDa FITC-dextran (Sigma-Aldrich) in the Chamber 2 collagen hydrogel. FITC-dextran was incorporated from a 350 µg mL□¹ stock solution to obtain a final concentration of 3.5 µg mL□¹, matching the initial concentration used for the VEGF-A diffusion studies. Chamber 2 was loaded on the day before imaging and allowed to polymerize overnight at 4 °C, while Chamber 1 was prepared using the same collagen I–fibrin hydrogel formulation described above. Fluorescence images were acquired using a Zeiss LSM800 confocal microscope under identical acquisition settings (laser power, detector gain and exposure time) at 4, 8, 12, 16, 20 and 24 h after hydrogel loading. Fluorescence intensity profiles were extracted in Fiji/ImageJ along predefined regions of interest encompassing Chamber 2, Channel 2, Reservoir 2, the microchannel array, Reservoir 1 and Chamber 1. Representative images were displayed using the Fire lookup table (LUT) to enhance visualization of the diffusion gradient, whereas quantitative fluorescence measurements were performed on the original fluorescence images. Experimental fluorescence profiles were subsequently compared with the COMSOL-predicted diffusion profiles.

#### Automated quantification of cell migration

Migrated cells were quantified using a custom image-analysis pipeline developed in FIJI/ImageJ. Brightfield images were used to automatically segment the microchannel array, generating a binary mask that defined the analysis region. Fluorescence images acquired from the corresponding field of view were then analyzed to identify individual fluorescent objects, which were filtered according to user-defined thresholds for projected area, circularity, fluorescence intensity, and overlap with the segmented microchannels. Only fluorescent objects satisfying all selection criteria were classified as migrated cells. The analysis generated both annotated overlay images for quality control and quantitative output files containing the number of migrated cells together with the morphological parameters of each detected object. Automated detections were visually verified and manually corrected only when necessary.

#### Statistical analysis

Cell counts obtained from independent microfluidic devices were analyzed using GraphPad Prism (GraphPad Software, San Diego, CA, USA). Data are presented as mean ± standard deviation (SD), with individual data points representing independent microfluidic devices. Differences among experimental groups were assessed using ordinary one-way ANOVA followed by Dunnett’s multiple-comparisons test against the corresponding control group. Statistical significance was defined as *P* < 0.05.

## Results

Heterogeneous expression of VEGF receptors across pediatric solid tumors supports the evaluation of VEGF-A165 and VEGF-C signaling

To identify clinically relevant chemotactic cues to evaluate in the MET-on-a-Chip platform, we first explored the expression of the main VEGF receptors across publicly available pediatric solid tumor datasets included in the R2 MegaSampler platform. Because VEGF-A165 and VEGF-C primarily signal through VEGFR2 (KDR) and VEGFR3 (FLT4), respectively, we analyzed the expression of both receptors across neuroblastoma, Ewing sarcoma and osteosarcoma, comparing primary lesions and metastatic samples whenever datasets were available.

KDR was broadly expressed across all three tumor types, although its expression varied considerably between independent datasets and between primary and metastatic lesions **(Figure 1A)**. In contrast, FLT4 showed a more homogeneous expression pattern across datasets and was consistently detected in all tumor types, with the highest expression levels observed in Ewing sarcoma and osteosarcoma **(Figure 1B)**. Despite the heterogeneity observed across independent patient cohorts, both VEGFR2 (KDR) and VEGFR3 (FLT4) were consistently represented in pediatric solid tumors. Also, both receptors were detected in datasets derived from primary tumors as well as metastatic lesions, indicating that VEGF signaling remains represented across different stages of disease progression. Based on these observations, VEGF-A165 and VEGF-C were selected as representative vascular and lymphatic chemotactic cues for subsequent evaluation in the MET-on-a-Chip platform.

**Figure 1.**
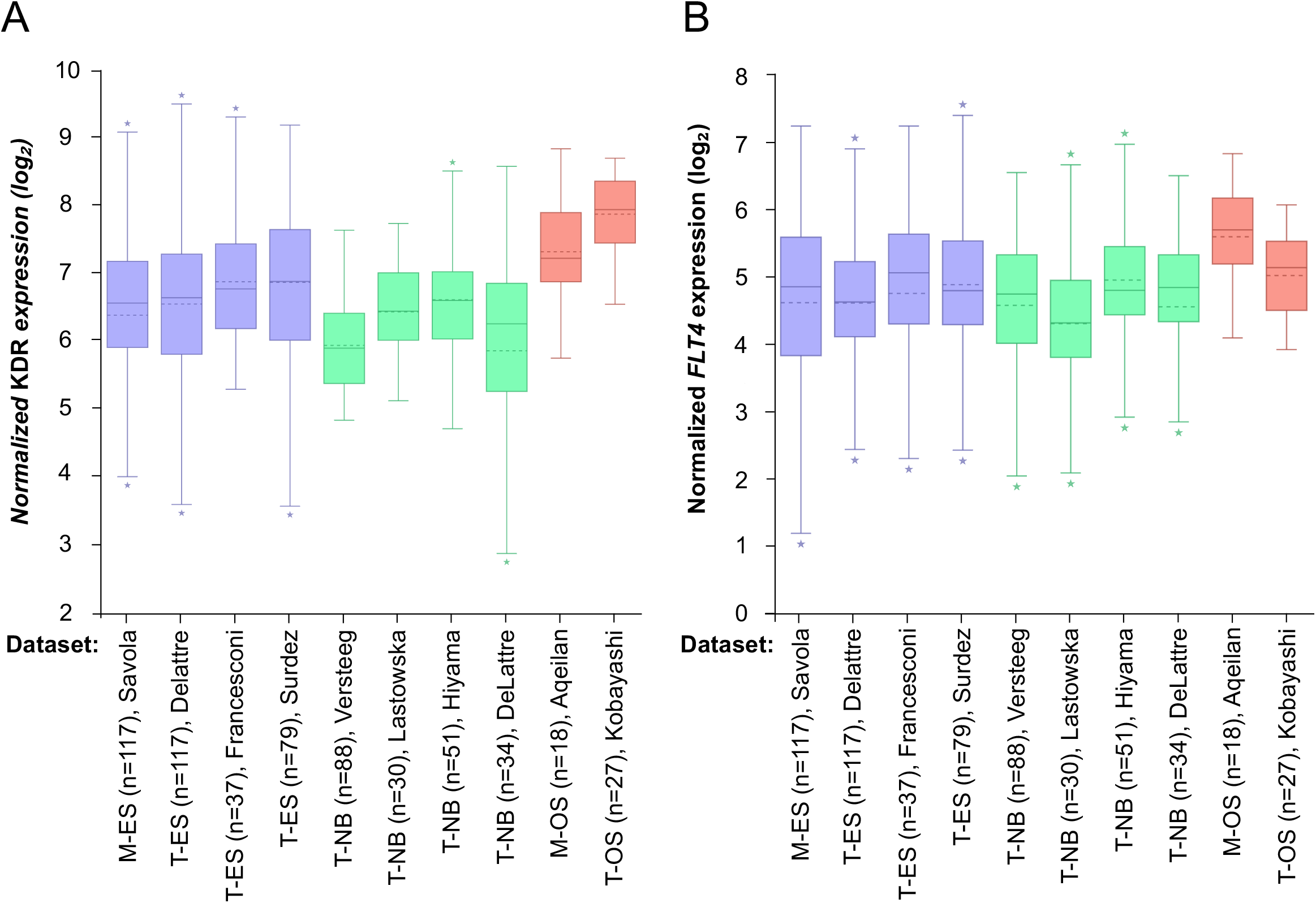
Heterogeneous expression of VEGF receptors across pediatric solid tumor datasets. **(A)** Expression of KDR, encoding VEGFR2, and **(B)** FLT4, encoding VEGFR3, across publicly available pediatric solid tumor datasets included in the R2 MegaSampler platform. Expression values are presented as log2-transformed signal intensity. The distributions illustrate inter-dataset and inter-tumor heterogeneity in the expression of receptors associated predominantly with VEGF-A and VEGF-C signaling, respectively. M, metastasis; T, primary tumor; ES, Ewing sarcoma; NB, neuroblastoma; OS, osteosarcoma.

### Development and fabrication of a minimally functional MET-on-a-Chip platform

To experimentally evaluate the chemotactic activity of VEGF-A165 and VEGF-C under controlled conditions, we developed a minimally functional metastasis-on-a-chip (MET-on-a-Chip) platform specifically designed to isolate the contribution of soluble gradients to tumor cell migration. Rather than reproducing the full complexity of the metastatic niche, the platform was intentionally designed as the simplest system capable of quantitatively evaluating the contribution of individual chemotactic cues to tumor cell migration.

The MET-on-a-Chip platform consists of two independent culture chambers (Chamber 1, Ch1, and Chamber 2, Ch2) connected to two intermediate reservoirs (Reservoir 1, R1, and Reservoir 2, R2) through 300-µm-wide access microchannels (Channels 1 and 2) (**Figure 2A**). The reservoirs are interconnected by an array of 125 parallel microchannels, which define the diffusion interface between both compartments. These migration microchannels were designed with a width of 15 µm to reproduce the physical confinement encountered by tumor cells during microvascular dissemination. Although smaller than the diameter of most tumor cells, this geometry remains permissive for cell migration while imposing physiologically relevant mechanical constraints. Their 1.38-mm length, together with the millimeter-scale separation between both culture chambers, establishes a long-range migration path while maintaining physical separation between the chemotactic source and the tumor compartment.

**Figure 2.**
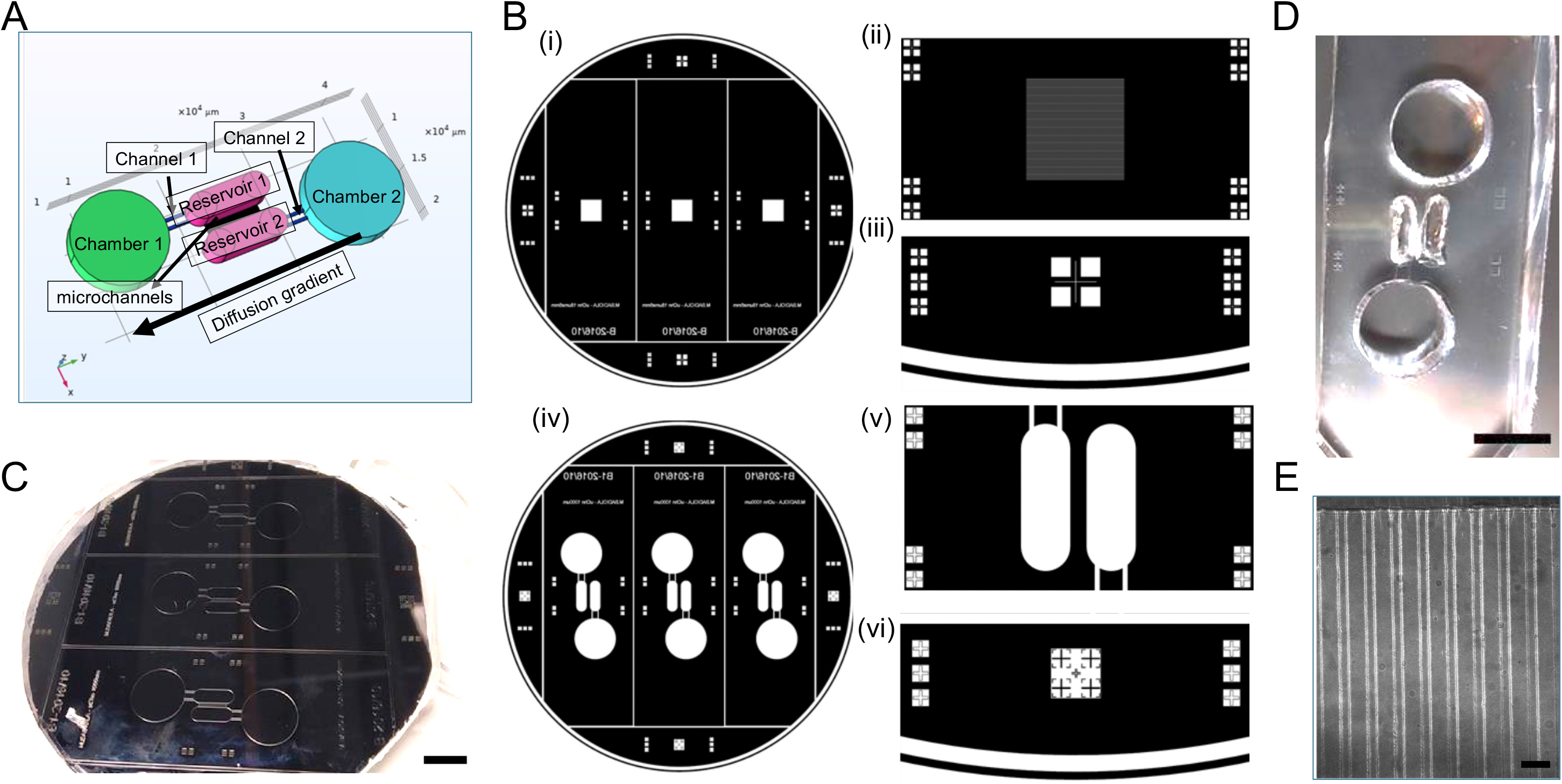
Design and fabrication of the MET-on-a-Chip platform for quantitative chemotaxis studies. **(A)** Three-dimensional representation of the MET-on-a-Chip device generated in COMSOL Multiphysics. The platform consists of two culture chambers connected through opposing reservoirs by an array of parallel microchannels. The predicted direction of chemoattractant diffusion from Chamber 2 toward Chamber 1 is indicated. (B) Acetate photomasks used to fabricate the two-layer SU-8 master mold. (i) First-layer mask, showing (ii) the microchannel array and (iii) the corresponding alignment marks. (iv) Second-layer mask, showing (v) the chambers and reservoirs separated by 1 mm and (vi) the corresponding alignment marks. **(C)** Representative SU-8 master fabricated on a 4-inch silicon wafer. Scale bar = 1 cm. **(D)** Representative PDMS replica of the MET-on-a-Chip device bonded to a glass coverslip. Scale bar = 1 cm. **(E)** Bright-field micrograph of the microchannel array connecting Reservoir 1 and Reservoir 2. Scale bar = 50 µm.

The device was fabricated by two-layer SU-8 photolithography followed by PDMS soft lithography (**Figure 2B–D**). The first lithographic layer defined the microchannel array, whereas the second generated the culture chambers, access microchannels, reservoirs and alignment features required for accurate layer registration. Optical microscopy confirmed the successful fabrication and integrity of the final microchannel array (**Figure 2E**).

Geometrical characterization demonstrated high fabrication reproducibility. The culture chambers measured 10.25 ± 0.21 mm (Ch1) and 10.82 ± 0.26 mm (Ch2) in diameter, whereas the intermediate reservoirs were approximately 3.6–3.7 mm wide and 9.2 mm long. The migration interface consisted of 125 parallel microchannels measuring 15.0 ± 0.2 µm in width, 1.38 ± 0.19 mm in length and separated by 34.9 ± 0.2 µm. In addition, interferometric measurements demonstrated that different PDMS curing protocols did not significantly affect microchannel depth, confirming the robustness of the fabrication process. The experimentally measured dimensions of the final device were subsequently used to reconstruct the computational geometry employed for all subsequent COMSOL simulations.

### COMSOL simulations identify the VEGF-A165 loading concentration and cell location required to generate biologically relevant exposure

Before performing the migration experiments, COMSOL simulations were used to rationally determine the VEGF-A165 loading concentration required to establish a stable concentration gradient across the MET-on-a-Chip platform. Experimentally, VEGF-A165 will be incorporated into a collagen hydrogel placed in Chamber 2, while tumor cells will be embedded in a separate hydrogel designed to mimic the tumor microenvironment. Because VEGF-A165 must diffuse from Chamber 2 through Channel 2, Reservoir 2, the microchannel array, and Reservoir 1 before reaching the opposite side of the device, its spatial and temporal distribution was modeled before selecting the final position of the cell-containing hydrogel.

Several initial VEGF-A165 loading concentrations were evaluated in silico (3.5 ng/mL, 1 µg/mL, and 3.5 µg/mL; additional simulations not shown). The objective was not to maximize VEGF-A165 concentration within the device, but to identify the lowest loading concentration capable of producing detectable and sustained exposure in potential cell-seeding regions over the duration of the migration assay. The selection was guided by reported VEGF-A concentrations in cancer patients, including approximately 13.3–39.4 pg/mL in tumor interstitial tissue, 74 pg/mL in blood, and 328 pg/mL in plasma^21^.

The two lower loading concentrations produced negligible VEGF-A165 levels on the opposite side of the platform and were not pursued experimentally. For the selected concentration of 3.5 µg/mL, time-dependent profiles were extracted along predefined 3D Cut Lines covering consecutive regions of the device (**Figure 3A–D**). VEGF-A165 concentrations were highest in Channel 2, immediately downstream of the collagen hydrogel, and decreased progressively upon entering Reservoir 2 and crossing the microchannel array. Concentrations increased over time in all receiving compartments, with profiles evaluated between 4 and 24 h.

**Figure 3.**
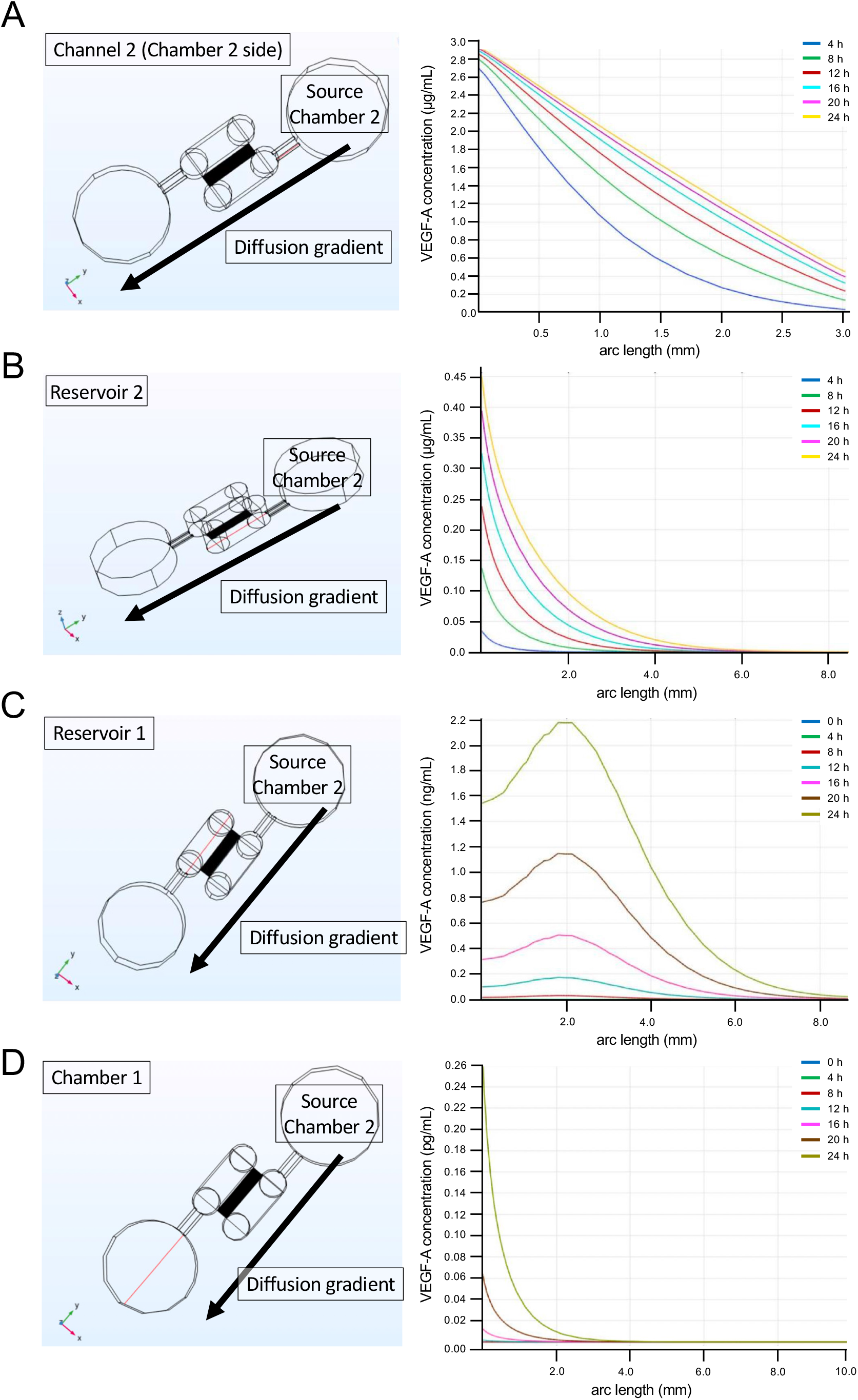
COMSOL prediction of spatial and temporal VEGF-A distribution within the MET-on-a-Chip device. VEGF-A transport was simulated following the incorporation of 3.5 µg/mL VEGF-A into Chamber 2. For each region, the left panel indicates the three-dimensional cut line used to extract the concentration profile, and the right panel shows the predicted VEGF-A concentration as a function of position and time. Profiles were obtained **(A)** along Channel 2, **(B)** within Reservoir 2, adjacent to the VEGF-A source, **(C)** in Reservoir 1, immediately beyond the microchannel array, and **(D)** in Chamber 1. The simulations predict progressive spatial attenuation of VEGF-A across the device, with high local concentrations near the source and substantially lower concentrations reaching Chamber 1. Different concentration units and y-axis ranges were used for the individual device regions.

After passage through the microchannel array, Reservoir 1 showed a spatial concentration maximum close to the microchannel outlet (**Figure 3C**). VEGF-A165 levels in this region increased from negligible concentrations during the first hours to the ng/mL range at later time points, reaching approximately 2.2 ng/mL at the local maximum after 24 h. This result showed that cells positioned in Reservoir 1 would be exposed to a clearly detectable VEGF-A165 gradient immediately after the growth factor crossed the microchannel barrier.

Further diffusion toward Chamber 1 resulted in an additional reduction of several orders of magnitude. At 24 h, the predicted concentration was approximately 17 pg/mL at the entrance of Channel 1, adjacent to Chamber 1, which falls within the reported range for tumor interstitial VEGF-A165 concentrations. Within Chamber 1 itself, however, the predicted concentration near the inlet was approximately 0.25 pg/mL after 24 h (**Figure 3D**), while concentrations at earlier time points remained substantially lower.

These simulations showed that the concentration incorporated into the collagen hydrogel cannot be interpreted as the concentration experienced by the cells. They also indicated that cell location within the platform strongly determines VEGF-A165 exposure. Loading Chamber 2 with 3.5 µg/mL VEGF-A165 generated biologically relevant concentrations near the entrance to Chamber 1, but substantially greater and earlier exposure in Reservoir 1. This condition was therefore selected for experimental validation, while the final cell-seeding configuration was defined based on the predicted concentration ranges in each compartment. Consequently, subsequent migration experiments were performed using Configuration 2, in which tumor cells were positioned in Reservoir 1 adjacent to the microchannel array to maximize exposure to the predicted gradients.

### COMSOL simulations identified both the VEGF-C loading concentration and the optimal region for cellular exposure

Because VEGF-C was expected to establish concentration gradients different from those predicted for VEGF-A165, an independent computational analysis was performed before the migration experiments. The objective was not only to determine the VEGF-C concentration required in the collagen hydrogel placed in Chamber 2, but also to identify the region of the microfluidic device where tumor cells would be exposed to biologically relevant concentrations during the experimental time window.

Two initial VEGF-C loading concentrations were evaluated in silico (0.1 and 5 µg/mL). The lower concentration generated negligible VEGF-C levels throughout the device over the 48-h simulation period and was therefore considered insufficient to establish a measurable chemotactic gradient (**Figure 4A–C**). In contrast, loading the collagen hydrogel with 5 µg/mL produced sustained VEGF-C transport across the platform and was selected for subsequent experimental validation.

**Figure 4.**
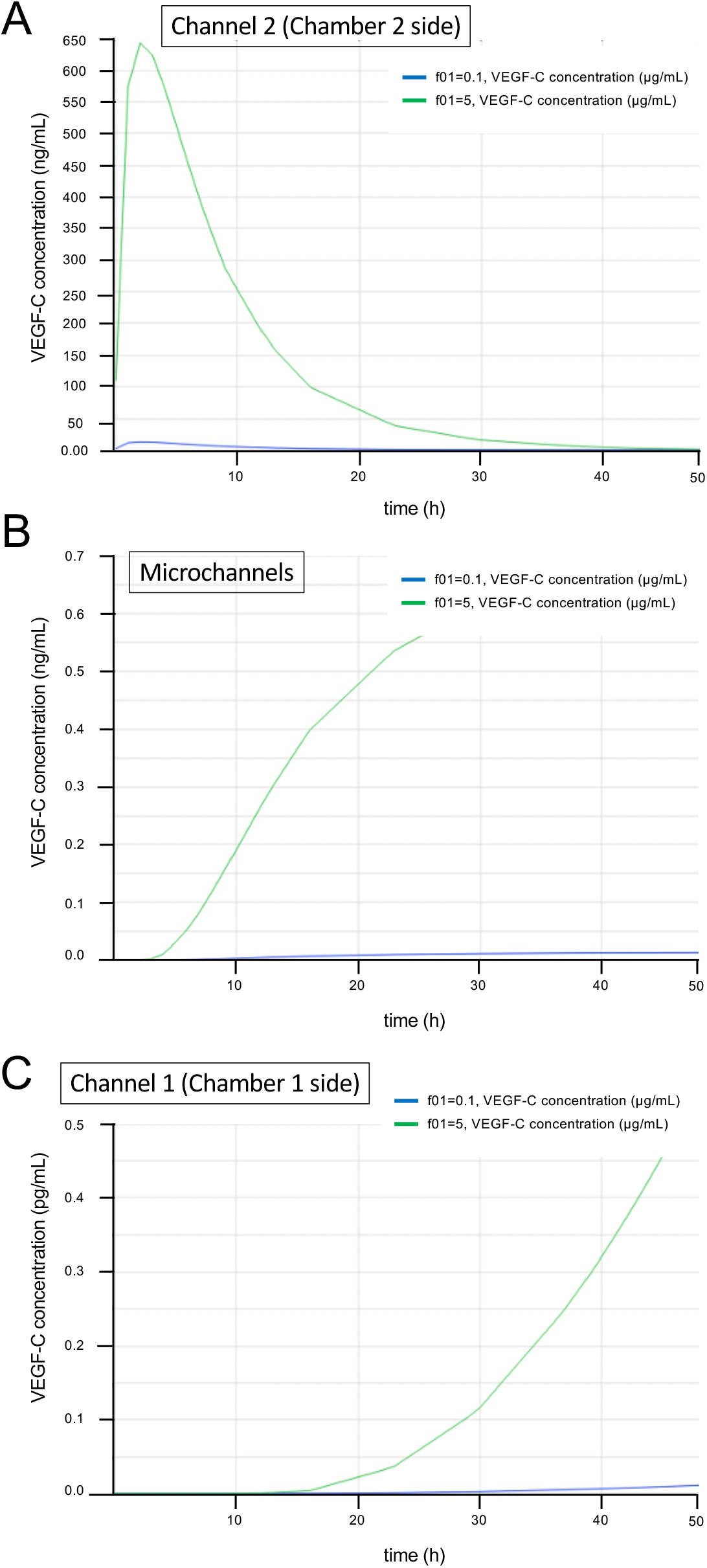
COMSOL-guided selection of the initial VEGF-C concentration and cellular exposure region. VEGF-C transport was simulated using initial concentrations of 0.1 or 5 µg/mL in Chamber 2. Predicted VEGF-C concentrations over time are shown in **(A)** Channel 2, adjacent to the source, **(B)** at the microchannel array separating Reservoirs 2 and 1, and **(C)** in Channel 1, on the Chamber 1 side of the device. An initial concentration of 5 µg/mL generated measurable exposure at the microchannel array within the experimental time window, whereas 0.1 µg/mL produced substantially lower concentrations and negligible predicted exposure at the microchannel array and in Channel 1. Different concentration units and y-axis ranges were used for the individual device regions.

Immediately adjacent to the source hydrogel (Channel 2), VEGF-C concentrations remained in the hundreds of ng/mL range during the first hours before progressively decreasing as diffusion proceeded (**Figure 4A**). As VEGF-C crossed the microchannel array, concentrations increased steadily over time, reaching approximately 0.5 ng/mL after 24 h (**Figure 4B**). Downstream of the microchannels, however, VEGF-C concentrations remained substantially lower, reaching only approximately 0.05 pg/mL at 24 h and approximately 0.45 pg/mL after 48 h at the entrance of Chamber 1 (**Figure 4C**).

These simulations demonstrated that, unlike VEGF-A165, the biologically relevant VEGF-C gradient was predicted to be concentrated around the microchannel array rather than within Chamber 1 during the experimental period. Consequently, migration experiments evaluating VEGF-C were designed by positioning tumor cells adjacent to Reservoir 1, immediately downstream of the microchannels, where the steepest and most sustained concentration gradients were predicted to occur.

### Experimental quantification of release of VEGF-A165 and VEGF-C from collagen hydrogels

To verify that the concentrations selected by the computational model were effectively released from the collagen hydrogels, VEGF-A165 and VEGF-C levels were quantified in the supernatant after 24 h under both hydrogel preparation (PBS, 4°C) and standard culture (RPMI, 37°C) conditions (**Table 1**).

**Table 1.**
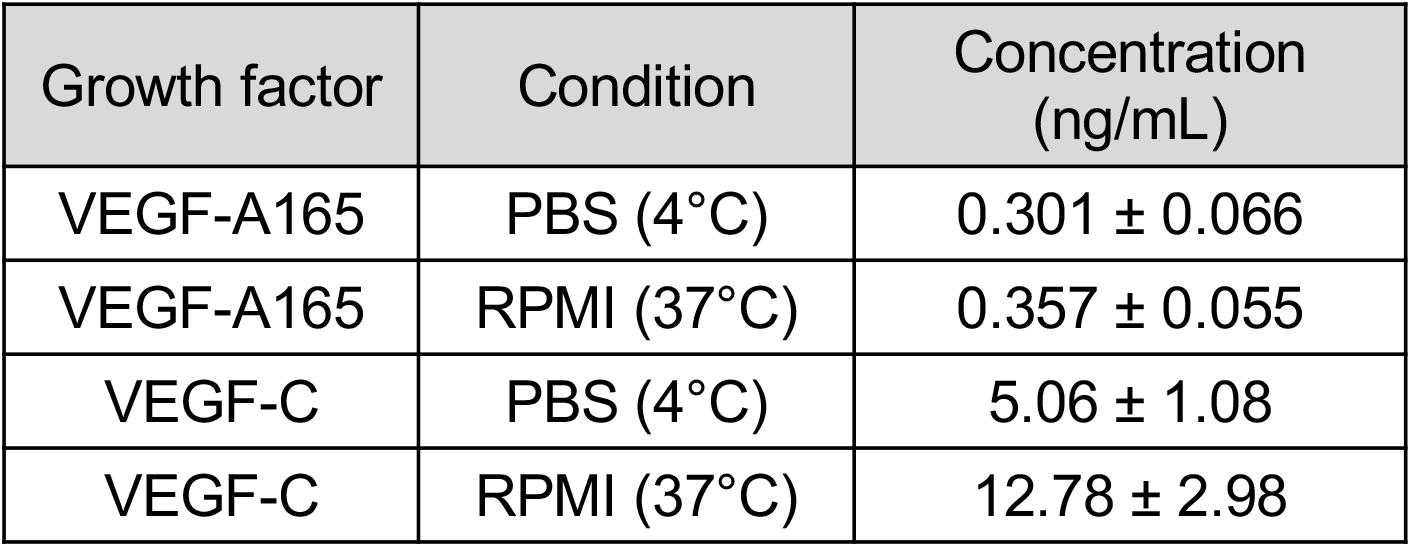
Quantification of VEGF release from collagen hydrogels loaded into Chamber 2 after 24 h. Concentrations of VEGF-A165 and VEGF-C released into the supernatant after 24 h of incubation under gel preparation conditions (PBS, 4°C) or standard cell culture conditions (RPMI, 37°C), as determined by ELISA. A total of 600 µL of supernatant was collected from each device. Data are presented as mean ± SD (n = 4 independent hydrogels per condition).

| Growth factor | Condition | Concentration (ng/mL) |
| --- | --- | --- |
| VEGF-A165 | PBS (4°C) | 0.301 ± 0.066 |
| VEGF-A165 | RPMI (37°C) | 0.357 ± 0.055 |
| VEGF-C | PBS (4°C) | 5.06 ± 1.08 |
| VEGF-C | RPMI (37°C) | 12.78 ± 2.98 |

VEGF-A165 release was comparable under both conditions, with concentrations of 0.301 ± 0.066 ng/mL in PBS and 0.357 ± 0.055 ng/mL in RPMI. In contrast, VEGF-C release was substantially higher, reaching 5.06 ± 1.08 ng/mL under polymerization conditions and increasing to 12.78 ± 2.98 ng/mL after incubation in complete culture medium.

These results demonstrate that both growth factors are released from the collagen matrix during the first 24 h, while also revealing distinct release profiles. The higher recovery observed for VEGF-C is consistent with its higher initial loading concentration, and confirms that sufficient soluble growth factor becomes available during the experimental time window used for the migration assays.

### Experimental validation of spatial diffusion across the microfluidic device

The spatial distribution of a 70-kDa fluorescent tracer was examined to determine whether macromolecules loaded into Chamber 2 could diffuse through the complete microfluidic pathway toward Chamber 1. The device layout and the direction of diffusion are shown in **Figure 5A**. FITC-dextran was introduced into the collagen hydrogel in Chamber 2, and fluorescence was evaluated sequentially in Channel 2, Reservoir 2, Reservoir 1, and Channel 1, following the expected diffusion path from the source toward the opposite side of the device.

**Figure 5.**
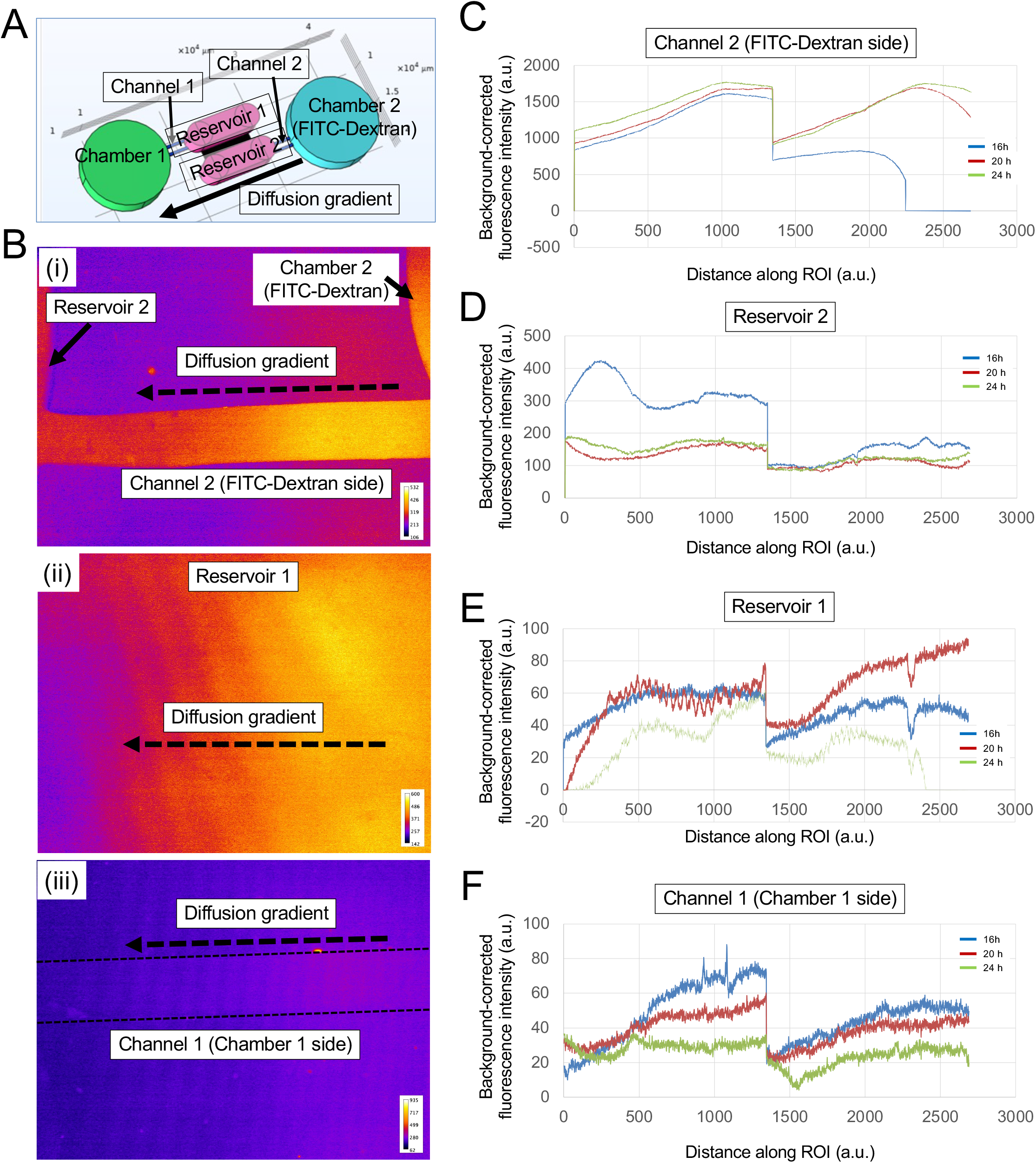
Experimental assessment of spatial diffusion within the microfluidic device using 70-kDa FITC-dextran. **(A)** Device geometry and direction of FITC-dextran diffusion from Chamber 2 toward Chamber 1. The analyzed regions comprised Channel 2, Reservoir 2, Reservoir 1, and Channel 1. **(B)** Representative fluorescence images acquired 16 h after loading FITC-dextran into Chamber 2, showing (i) Reservoir 2 and Channel 2 adjacent to the source, (ii) Reservoir 1 after diffusion across the microchannel array, and (iii) Channel 1 on the Chamber 1 side. Images are displayed using the Fire lookup table, with warmer colors indicating higher fluorescence intensity. Dashed arrows indicate the direction of diffusion. **(C–F)** Fluorescence-intensity profiles measured along predefined regions of interest (ROI) in **(C)** Channel 2, **(D)** Reservoir 2, **(E)** Reservoir 1, and (F) Channel 1 at 16, 20, and 24 h. Background-corrected fluorescence intensity is expressed in arbitrary units.

Representative fluorescence images acquired 16 h after FITC-dextran loading showed a clear spatial redistribution of the tracer across the device (**Figure 5B**). In the region immediately adjacent to the source, Chamber 2 and Channel 2 displayed the highest fluorescence signal, whereas the signal decreased toward Reservoir 2, generating a visible fluorescence gradient along the direction of diffusion (**Figure 5B-i**). FITC-dextran was also detected in Reservoir 1 after passage through the microchannel array. Within this region, fluorescence remained spatially heterogeneous, with higher intensities on the side closer to the microchannels and a progressive reduction toward Chamber 1 (**Figure 5B-ii**). A weaker but detectable signal was observed in Channel 1, confirming that the tracer had reached the distal side of the device while retaining a spatial intensity gradient (**Figure 5B-iii**).

Fluorescence-intensity profiles were subsequently quantified at 16, 20, and 24 h. Channel 2 consistently showed the highest fluorescence levels among all analyzed regions, with intensities reaching approximately 1,500–1,800 a.u. at 20 and 24 h (Figure 5C). The profiles increased along the analyzed region adjacent to Chamber 2, followed by a sharp reduction at the transition between spatial segments of the ROI. In the distal portion of Channel 2, fluorescence again increased with distance at 20 and 24 h, whereas the 16-h profile remained lower and declined to background levels near the end of the analyzed region.

In Reservoir 2, fluorescence intensities were substantially lower than in Channel 2 (**Figure 5D**). At 16 h, the profile reached approximately 400 a.u. in the region closest to the source and remained above the corresponding 20-and 24-h profiles across most of the first ROI segment. After the central transition in the profile, intensities decreased to approximately 80–180 a.u., with relatively similar values at the three time points. These measurements indicate a marked reduction in tracer concentration between Channel 2 and Reservoir 2.

Further from the source, FITC-dextran remained detectable in Reservoir 1 (**Figure 5E**). Fluorescence intensities were lower than in Reservoir 2 and varied with both position and time. At 20 h, the signal increased along the analyzed region and reached approximately 90 a.u. at its distal end. The 16-h profile showed intermediate values, generally ranging from approximately 30 to 60 a.u., whereas the 24-h profile remained lower and decreased sharply near the end of the ROI. Despite these temporal differences, the presence of fluorescence throughout Reservoir 1 demonstrated transport of the tracer across the microchannel array.

The lowest fluorescence intensities were measured in Channel 1, on the Chamber 1 side of the device (**Figure 5F**). At 16 h, fluorescence progressively increased along the first analyzed segment and reached approximately 70–80 a.u., while the corresponding 20-h profile reached approximately 50–60 a.u. Signal was still detectable at 24 h, although at lower levels. A similar ordering of the three time points was observed in the second segment of the ROI, with the 16-h profile remaining highest and the 24-h profile lowest. Overall, the progressive reduction in fluorescence intensity from Channel 2 to Reservoir 2, Reservoir 1, and Channel 1 confirmed that FITC-dextran crossed the microchannel array and reached the opposite side of the device. The spatially resolved profiles therefore experimentally support the formation of a macromolecular concentration gradient across the full MET-on-a-Chip geometry.

### VEGF-C selectively promotes migration of Ewing sarcoma and osteosarcoma cells through the microchannel array

To determine whether VEGF gradients promote tumor cell migration within the MET-on-a-Chip platform, neuroblastoma, Ewing sarcoma and osteosarcoma cells were cultured under control, VEGF-A165 and VEGF-C conditions, and the number of cells entering the microchannel array was quantified using the automated image-analysis pipeline described above.

Representative brightfield and fluorescence overlay images showed clear differences in the number of fluorescent cells detected within the microchannels depending on both tumor type and growth factor (**Figure 6A**). Neuroblastoma cells were only occasionally detected within the microchannel array under all three conditions, with no obvious increase in the presence of either VEGF-A165 or VEGF-C. In contrast, Ewing sarcoma cells exhibited a marked increase in the number of cells occupying the microchannels in response to VEGF-C, whereas only a modest increase was observed following VEGF-A□□□ treatment. A similar trend was observed for osteosarcoma cells, which showed a visibly higher number of migrated cells under VEGF-C compared with both the control and VEGF-A165 conditions.

**Figure 6.**
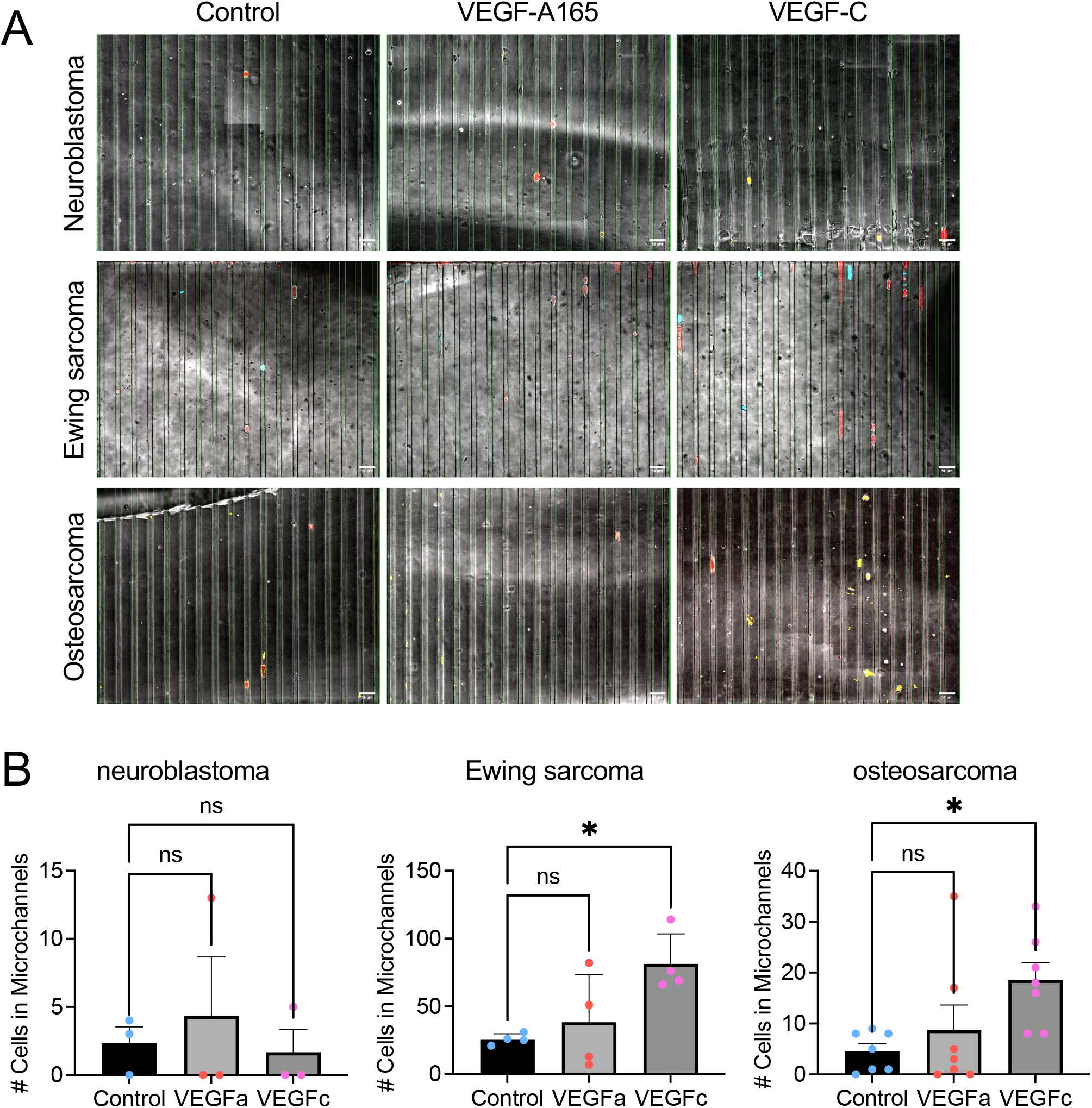
VEGF-C promotes tumor-type-specific migration through the MET-on-a-Chip microchannel array. **(A)** Representative merged bright-field and fluorescence images of neuroblastoma, Ewing sarcoma, and osteosarcoma cells detected within the microchannel array under control, VEGF-A165, and VEGF-C conditions. Images were processed using a custom Fiji macro that automatically identified the microchannel region, detected fluorescent cells, and retained cells located within the microchannels according to predefined size, circularity, intensity, and channel-overlap criteria. Scale bars =50µm. **(B)** Quantification of the number of cells detected within the microchannels for neuroblastoma, Ewing sarcoma, and osteosarcoma. Individual data points represent independent microfluidic devices, and bars show mean ± SD. Statistical significance was assessed using ordinary one-way ANOVA followed by Dunnett’s multiple-comparisons test against the corresponding control. *p < 0.05; ns, not significant.

Quantification of migrated cells confirmed these observations (**Figure 6B**). Neuroblastoma cells showed very limited migration into the microchannel array irrespective of the experimental condition, and neither VEGF-A165 nor VEGF-C induced a statistically significant increase compared with the control. In contrast, VEGF-C significantly increased the number of Ewing sarcoma cells detected within the microchannels compared with untreated controls (*P* < 0.05), whereas VEGF-A165 did not reach statistical significance. Likewise, osteosarcoma cells responded significantly to VEGF-C but not to VEGF-A165, demonstrating a selective pro-migratory effect of VEGF-C in these tumor types.

Overall, these results indicate that VEGF-C, but not VEGF-A165, acts as an effective chemoattractant for Ewing sarcoma and osteosarcoma cells within the MET-on-a-Chip platform, whereas neuroblastoma cells exhibited minimal migration under either growth factor condition.

## Discussion

In this study, we developed a computationally guided metastasis-on-a-chip platform to determine whether individual soluble cues are sufficient to direct the migration of pediatric cancer cells through a spatially confined microchannel array. Rather than attempting to reproduce the full complexity of the metastatic niche, the platform was intentionally designed according to the concept of the Minimally Functional Unit (MFU), incorporating only those biological and engineering elements required to answer a single experimental question under controlled conditions. By combining predictive computational modeling with experimental validation, the present workflow establishes a rational strategy for designing metastasis-on-a-chip assays in which device geometry, molecular transport, growth factor loading, and cell positioning are defined before biological experimentation. Using this approach, we demonstrate that both experimental configuration and the identity of the chemotactic cue critically determine whether a measurable migratory phenotype can be resolved.

One of the main methodological insights provided by this study is that the concentration initially loaded into a microfluidic device should not be interpreted as the concentration ultimately experienced by the cells. Although VEGF-A165 and VEGF-C were incorporated into the collagen hydrogel at concentrations in the microgram-per-milliliter range, computational modeling predicted a progressive attenuation of both ligands as they diffused through the connecting channel, intermediate reservoirs, and microchannel array. Consequently, local exposure differed by several orders of magnitude depending on the position of the responding cells within the device. This observation has important implications beyond the present study. In many microfluidic migration assays, cells are positioned according to geometrical convenience rather than quantitative prediction of ligand exposure. Our results demonstrate that relatively small differences in compartment organization can determine whether cells are exposed to a biologically meaningful gradient^21^ during the experimental window and, therefore, whether directional migration can be detected at all.

This finding also illustrates the value of integrating computational modeling into assay design rather than using it only to interpret experimental observations retrospectively. Microfluidic systems offer precise spatial control over the cellular microenvironment, yet their behavior depends on multiple interacting variables, including geometry, molecular diffusion, matrix properties, transport distance, and cell location^13,14^. Optimizing these parameters exclusively through experimental iteration requires repeated fabrication cycles and extensive empirical testing, consuming time, reagents, and biological material before informative conditions are achieved. Here, finite-element simulations allowed us to predict concentration profiles throughout the device, define the growth factor loading required to achieve physiologically relevant local exposure, and identify the optimal region for cell seeding before any migration experiments were performed. In this context, COMSOL modeling becomes an integral component of experimental design rather than a simple engineering exercise.

More broadly, we believe that this workflow represents an evolution in the design of human microphysiological systems. More than a decade ago, we proposed the concept of the Minimally Functional Unit (MFU) as the simplest experimental system capable of reproducing the biological function required to answer a specific scientific question^15^. The underlying principle was that biological complexity should only be incorporated when it contributes mechanistic information that cannot be obtained using a simpler model. The present work extends this concept beyond the selection of biological components. Our results demonstrate that the functionality of a microphysiological system is determined not only by the cells and extracellular matrix that it contains, but also by its physical architecture, molecular transport properties, and the local concentrations ultimately experienced by the cells. From this perspective, predictive computational modeling becomes part of the MFU itself, allowing biological complexity and engineering design to be optimized simultaneously before experimental validation.

The experimental measurements support this computational strategy while also illustrating the complementary roles of predictive modeling and biological validation. FITC-dextran diffusion confirmed that macromolecules diffuse throughout the complete microfluidic pathway and establish long-range concentration gradients across the device. Likewise, ELISA measurements demonstrated the progressive release of both VEGF-A165 and VEGF-C from collagen hydrogels during the experimental period. Although inert dextran molecules cannot reproduce the complex matrix interactions, degradation kinetics, or receptor binding properties of VEGF family members, these experiments collectively validate the fundamental transport assumptions underlying the computational model. Rather than attempting to measure local ligand concentration directly throughout the device (a technically challenging task), the combination of predictive simulations with independent experimental validation provides a practical framework for designing and interpreting diffusion-based microphysiological systems. Importantly, the computational simulations also revealed that VEGF-A165 and VEGF-C required different loading concentrations to generate comparable local exposure within the cellular compartment. Despite these different initial loading conditions, both ligands ultimately reached the responding cells within a similar order of magnitude after diffusion across the complete device. This observation substantially reduces the possibility that the distinct biological responses observed in the migration assays simply reflect differences in ligand availability or transport efficiency. Instead, once molecular transport was normalized through computational design, the remaining differences are more plausibly explained by intrinsic biological responses of the tumor cells to vascular and lymphatic VEGF-family signaling. In this sense, computational modeling did not merely improve the experimental workflow; it enabled the biological comparison itself by controlling one of the major sources of experimental variability in diffusion-based microfluidic assays.

This study also highlights an important aspect of organ-on-a-chip development. Increasing biological complexity is often considered equivalent to improving physiological relevance, but more complex models also introduce additional variables that can make mechanistic interpretation more difficult. The objective of an experimental model is not to recreate the entire human body in vitro, but to isolate and quantitatively study a defined biological function under controlled conditions. The concept of the MFU provides a framework for this approach, in which complexity is introduced progressively and only when a simpler system is no longer sufficient to answer the biological question.

Although demonstrated here using VEGF-A165 and VEGF-C, the same workflow can be adapted to other soluble factors, cytokines, chemokines, extracellular vesicles, or combinations of signals involved in metastatic dissemination. Additional components, including endothelial or lymphatic barriers, stromal cells, immune populations, organ-specific cells, or extracellular matrices, can then be incorporated one at a time when required by the scientific question, while maintaining experimental control and allowing the contribution of each component to be evaluated separately.

One of the most interesting observations emerging from this study is that controlling molecular transport computationally allowed the biological contribution of individual VEGF family members to be examined under comparable experimental conditions. Although VEGF-A165 and VEGF-C required different loading concentrations because of differences in the predicted concentration profiles, both ligands ultimately reached the cellular compartment within a similar order of magnitude after diffusion across the device. Consequently, the markedly different migratory responses observed cannot be readily attributed to differences in diffusion or ligand availability alone. Instead, they reveal intrinsic differences in how pediatric tumor cells interpret vascular and lymphatic VEGF-family signals.

Under these conditions, VEGF-C consistently promoted migration of both Ewing sarcoma and osteosarcoma cells through the confined microchannel array, whereas VEGF-A165 failed to induce a comparable response in any of the three pediatric tumor models investigated. Importantly, migration in the present platform required considerably more than simple cell displacement. Cells were required to detect the soluble gradient, polarize, deform, and actively migrate through 15-µm-wide confined microchannels before entering the opposite compartment. The observed phenotype therefore represents an integrated functional response involving chemotactic sensing together with the ability to migrate under physical confinement, two processes that are both essential during metastatic dissemination.

The lack of response to VEGF-A165 deserves particular consideration because VEGF-A has long been associated with tumor progression and metastasis across multiple cancer types. However, the majority of studies supporting this association describe indirect biological functions of VEGF-A rather than direct chemotactic effects on tumor cells^18,19^. VEGF-A is a master regulator of angiogenesis and vascular permeability, promotes endothelial cell proliferation and survival, remodels the extracellular matrix through vascular activation, and influences stromal and immune cell recruitment. Many of its pro-metastatic effects therefore emerge from coordinated interactions between tumor cells and the surrounding microenvironment rather than from direct stimulation of tumor-cell motility^18,19^. The present MET-on-a-Chip was intentionally designed to exclude endothelial, stromal, and immune compartments in order to isolate the contribution of soluble factors acting directly on tumor cells. Consequently, the absence of a detectable migratory response to VEGF-A165 should not be interpreted as evidence that VEGF-A165 lacks biological relevance in pediatric tumors. Rather, our results indicate that VEGF-A165 alone is insufficient to drive directional migration under these experimentally controlled conditions and suggest that additional microenvironmental components are required to fully reproduce its biological activity.

In contrast, VEGF-C retained promigratory activity even within this highly simplified system. VEGF-C is best known for its role in lymphangiogenesis through activation of VEGFR3, although proteolytically processed VEGF-C can also activate VEGFR2 depending on its maturation state^22^. Increasing evidence suggests that, beyond its effects on lymphatic endothelial cells, VEGF-C may directly regulate the behavior of malignant cells in selected tumor types. Our findings support this possibility in pediatric sarcoma models by demonstrating that VEGF-C alone is sufficient to increase migration in the absence of endothelial or lymphatic compartments. While the receptor responsible for this phenotype remains to be experimentally identified, these results provide a strong rationale for future studies combining receptor expression analysis with pharmacological or genetic inhibition of VEGFR2 and VEGFR3.

The response observed in Ewing sarcoma is particularly noteworthy because VEGF signaling in this disease has traditionally been investigated from the perspective of tumor angiogenesis and vascularization. Previous studies have shown that VEGF-A contributes to vascular development within Ewing sarcoma tumors and that inhibition of VEGF signaling reduces tumor vascularization and growth in vivo. Recent clinical data suggest that these results may also extend to patients^23,24^. Our results do not contradict these observations. Instead, they separate two distinct biological functions that are frequently considered together. A signaling pathway may contribute substantially to metastatic progression through its effects on endothelial cells and vascular remodeling while having little or no direct chemotactic activity on tumor cells themselves. By intentionally removing endothelial and stromal components, the present platform demonstrates that VEGF-C, rather than VEGF-A165 functions as the dominant soluble migratory cue for SK-N-MC cells under the conditions tested.

A similar interpretation applies to osteosarcoma. VEGF-A expression has repeatedly been correlated with angiogenesis, pulmonary metastasis, and poor clinical outcome^25^, providing a strong rationale for anti-angiogenic therapeutic approaches. Nevertheless, these clinical associations do not necessarily imply that soluble VEGF-A directly guides tumor-cell migration. In the present study, U2OS cells responded robustly to VEGF-C but not to VEGF-A165, suggesting that these ligands contribute differently to metastatic progression. While VEGF-A165 may primarily influence the vascular microenvironment, VEGF-C appears capable of acting directly on tumor cells to promote migration under confined conditions. Although these observations currently derive from a single osteosarcoma cell line, they identify VEGF-C as an underexplored candidate regulator of osteosarcoma dissemination that warrants further mechanistic investigation.

In contrast, SK-N-BE(2) neuroblastoma cells displayed minimal migration under basal conditions and remained largely unresponsive to either VEGF-A165 or VEGF-C. Importantly, this negative result is biologically informative rather than simply inconclusive. It demonstrates that the MET-on-a-Chip does not produce a generalized increase in migration in response to any VEGF ligand and highlights the selective nature of the assay. Neuroblastoma metastasis, particularly to bone marrow, is known to depend on complex interactions involving extracellular matrix composition, stromal cells, chemokines, developmental cell state, and tissue-specific microenvironmental signals^26,27^. The absence of a response to isolated VEGF gradients therefore suggests that neither VEGF-A165 nor VEGF-C alone is sufficient to reproduce these processes in SK-N-BE(2) cells.

Finally, the comparison with the transcriptomic analysis presented in Figure 1 further emphasizes the importance of experimentally validating computational and omics-derived hypotheses. The heterogeneous expression of KDR and FLT4 across pediatric patient datasets provided the rationale for investigating VEGF-family signaling, but bulk transcriptomic data cannot identify the cellular origin of these receptors or predict the functional behavior of individual tumor models. Endothelial cells, stromal populations, and malignant cells all contribute to bulk tumor expression profiles. Consequently, transcript abundance alone cannot establish whether a particular tumor cell population expresses functional receptor protein or responds to ligand stimulation. By integrating transcriptomic analyses with computational modeling and functional migration assays, the present workflow illustrates how patient-derived molecular information can be translated into experimentally testable biological hypotheses within controlled human microphysiological systems.

Several limitations define the scope of the present study and naturally identify the next experimental steps. First, only one representative cell line from each pediatric tumor type was evaluated. Although these models capture distinct biological behaviors, they cannot represent the full heterogeneity of neuroblastoma, Ewing sarcoma, or osteosarcoma. Expanding the platform to additional cell lines, patient-derived tumor cells, organoids, or primary biopsy material will be essential to determine whether these findings are consistent across different tumor models, and to identify tumor-specific patterns of VEGF sensitivity.

Second, the current assay quantified migration as the number of cells entering the confined microchannel array after a defined experimental time point. Although this endpoint provides a robust and reproducible measure of directional migration under confinement, it does not distinguish between changes in migration speed, persistence, polarity, or the proportion of cells capable of initiating migration. Integration of biosensors would allow continuous monitoring of cell behavior, and provide a more comprehensive description of the migratory process. Alternatively, automated fluorescence microscopy combined with time-lapse imaging could provide continuous visualization of multiple chips simultaneously without requiring any modification of the current platform, representing a practical solution for laboratories without biosensor fabrication capabilities. Likewise, pharmacological inhibition or genetic manipulation of VEGFR2 and VEGFR3 will be required to determine the molecular mechanisms responsible for the VEGF-C-dependent phenotype identified here.

In the longer term, this approach may also facilitate the development of personalized metastasis-on-a-chip models. Incorporating patient-derived tumor cells or organoids into computationally optimized microphysiological platforms could allow individual metastatic behaviors to be investigated under human-relevant conditions while systematically evaluating candidate anti-metastatic therapies. Rather than serving exclusively as disease models, these systems could become predictive preclinical platforms for prioritizing therapeutic strategies before evaluation in more complex animal models or clinical studies. More broadly, combining predictive computational design with experimentally validated human microphysiological systems represents a promising strategy for improving reproducibility, reducing experimental trial-and-error, and accelerating the translation of engineered models into clinically relevant applications.

## Conclusions

We developed a computationally guided MET-on-a-Chip platform that combines predictive finite-element modeling with experimental validation to investigate cue-directed migration in pediatric solid tumors under quantitatively controlled conditions. By integrating transport simulations into the experimental design process, the platform rationally defines growth factor loading, cell positioning, and device configuration before biological experimentation, substantially reducing empirical optimization while improving mechanistic interpretation.

Using this workflow, we demonstrate that VEGF-C, but not VEGF-A165, is sufficient to promote directional migration of Ewing sarcoma and osteosarcoma cells through confined microchannels, whereas neuroblastoma cells remain largely unresponsive under identical conditions. These findings emphasize that molecular identity alone does not determine migratory behavior; rather, biologically meaningful responses emerge from the interaction between tumor-specific biology, local ligand exposure, and microphysiological system design.

More broadly, this study illustrates how predictive computational modeling can transform the development of metastasis-on-a-chip platforms from an empirical, trial-and-error process into a rational engineering workflow. Although demonstrated here using VEGF-family signaling as a proof of concept, the same strategy can be readily extended to other biological questions, providing a flexible and scalable framework for constructing minimally functional human microphysiological systems in which complexity is introduced only when required by the scientific question. Ultimately, we envision that this design philosophy will facilitate the development of more predictive experimental models capable of accelerating both mechanistic studies of metastasis and the preclinical evaluation of personalized anti-metastatic therapies.

## AUTHOR CONTRIBUTIONS

CM, LJ, AC, and EC performed experiments and contributed to data analysis and interpretation. MB and DF performed experiments. NGJ contributed to figure preparation. JM and JS reviewed the manuscript. AV conceived the project, performed experiments, analyzed and interpreted results, and wrote the manuscript.

## AVAILABILITY OF DATA AND MATERIALS

The datasets generated and/or analysed during the current study are available from the corresponding author upon reasonable request.

## COMPETING INTERESTS

The authors declare that they have no competing interests.

## ACKNOWLEDGMENTS

The IBEC Group is supported by Biomedical Research Networking (CIBER), Spain. CIBER is an initiative funded by the VI National R&D&i Plan 2008-2011, Iniciativa Ingenio 2010, Consolider Program, CIBER Actions, and the Instituto de Salud Carlos III (RD16/0006/0012; RD16/0011/0022), with the support of the European Regional Development Fund (ERDF). This work was funded by the CERCA Program and by the Commission for Universities and Research of the Department of Innovation, Universities, and Enterprise of the Generalitat de Catalunya (2017-SGR-1079).

AV is supported by grant INVES234968VILL from the Scientific Foundation of the Spanish Association Against Cancer, by grant CNS2023-143930 funded by MCIN/AEI/10.13039/501100011033 and the European Union NextGeneration EU/PRTR, by a grant from the Alternatives Research & Development Foundation (ARDF), 2025 Annual Open Grant Cycle (Application #1488946), and by the Association of Families and Friends of Children with Neuroblastoma (NEN).

We thank the MicroFabSpace and Microscopy Characterization Facility, Unit 7 of ICTS “NANBIOSIS” from CIBER-BBN at IBEC.

## Notes

### Competing Interest Statement

The authors have declared no competing interest.

